# Near-infrared spectroscopy outperforms genomics for predicting sugarcane feedstock quality traits

**DOI:** 10.1101/2020.07.16.206110

**Authors:** Mateus Teles Vital Gonçalves, Gota Morota, Paulo Mafra de Almeida Costa, Pedro Marcus Pereira Vidigal, Marcio Henrique Pereira Barbosa, Luiz Alexandre Peternelli

**Affiliations:** Departamento de Estatística, Universidade Federal de Viçosa, Viçosa, MG, Brazil; Department of Animal and Poultry Sciences, Virginia Polytechnic Institute and State University, Blacksburg, VA, United States of America; Instituto Federal Catarinense - Campus Concórdia, Concórdia, SC, Brazil; Centro de Análises de Biomoléculas/NuBioMol, Universidade Federal de Viçosa, Viçosa, MG, Brazil; Departamento de Fitotecnia, Universidade Federal de Viçosa, Viçosa, MG, Brazil

**Author notes:** (LAP).

## Abstract

The main objectives of this study were to evaluate the prediction performance of genomic and near-infrared spectroscopy (NIR) data and whether the integration of genomic and NIR predictor variables can increase the prediction accuracy of two feedstock quality traits (fiber and sucrose content) in a sugarcane population (*Saccharum* spp.). The following three modeling strategies were compared: M1 (genome-based prediction), M2 (NIR-based prediction), and M3 (integration of genomics s and NIR wavenumbers). Data were collected from a commercial population comprised of three hundred and eighty-five individuals, genotyped for single nucleotide polymorphisms (SNPs) and screened using NIR spectroscopy. We compared partial least squares (PLS) and BayesB regression methods to estimate marker and wavenumber effects. In order to assess model performance, we employed random sub-sampling cross-validation to calculate the mean Pearson correlation coefficient between observed and predicted genotypic values. Our results showed that models fitted using BayesB were most predictive than PLS models. We found that NIR (M2) provided the highest prediction accuracy, whereas genomics (M1) presented the lowest predictive ability, regardless of the measured traits and regression methods used. The integration of predictors derived from NIR spectroscopy and genomics into a single model (M3) did not significantly improve the prediction accuracy for the two traits evaluated. These findings suggest that NIR-based prediction can be an effective strategy for predicting the genotypic value of sugarcane clones.

## Introduction

The strides achieved with improved instruments, laboratory techniques, and bioinformatics tools have allowed the emergence of next-generation sequencing technologies [1]. These technologies can deliver DNA-level information at an ever more cost-effective and high-throughput manner and has boosted the important role genomic selection (GS) might play in plant breeding [2]. The idea of GS is to fit a regression model using phenotypic records and the entire set of molecular markers concurrently. The model developed enables the prediction of the genetic merit of genotyped but non-phenotyped populations [3].

The application of GS as a breeding strategy is envisioned to reduce costs while saving time and resources [4]. One example is the reduction of generation interval as genitors could be crossed, the resulting progeny have their DNA collected from seeds or juvenile tissues, and then genotyped to have their breeding values predicted, what may result in gains per unit of time [5]. Another advantage of GS is that it could eventually lead to a reduction in phenotyping costs, especially in situations where traits are difficult to measure and when there are thousands of candidate genotypes to evaluate [6]. Therefore, the adoption of GS over phenotypic selection is expected to augment selection efficiency and to accelerate cultivar release [7,8].

Nevertheless, the implementation of GS is highly dependent on accurate phenotyping records [9]. Moreover, conventional phenotyping is not excluded from GS-based plant breeding schemes as model updates on new training populations would still be necessary [10]. However, besides the forecasts of continuous advances and decreasing costs in molecular breeding, the phenotyping step remains a significant bottleneck [11]. Phenotyping routine traits at breeding stations are commonly performed manually over several crop years and across different environments, which is often a time-consuming, labor-intensive, and high-cost task [12]. Also, conventional phenotyping is prone to human error. Thus, the genetic potential of populations may not be fully exploited [11].

Research efforts to address these constraints are currently being conducted with the development of high-throughput phenotyping (HTP) systems. HTP is an incipient, though a growing area of interest among plant breeders [13]. These novel approaches include a multitude of sensors and imaging techniques mounted on ground-based or unmanned aerial vehicles that can collect phenotypes in a precise, automated, and large-scale fashion [14]. Thus, HTP tools have the potential to improve plant breeding program pipelines significantly [15]. Moreover, HTP technologies can replace standard less effective phenotyping protocols, thus saving much time and resources. For instance, near-infrared (NIR) spectroscopy technology has been successfully applied in agriculture to screen biological sample compositions and also for breeding purposes [9,16,17].

The combination of HTP systems with improved genomic tools is heralded to increase genetic gains in plant breeding [12,18]. However, the startling amount of high-throughput data being generated is outpacing our ability to explore it. Besides, how this information can properly be implemented is still unclear and needs further investigations [11,14]. The integration of HTP information into GS can be performed by the exploitation of HTP platforms to provide phenotypic records that can be either treated as secondary traits such as vegetation index or canopy temperature (correlated with core traits, e.g., yield) and regressed on molecular markers or as predictor variables together with molecular markers in a single- or multi-trait analysis [19]. Other strategies provide modeling alternatives that include interaction effects [20,21]. For instance, Crain et al. (2018) [22] investigated different proposals to integrate HTP derived variables into GS models in wheat and found improved prediction accuracies.

The main goal of this study was to investigate the performance of the integration of HTP and genomic datasets aiming to increase the accuracy of prediction for two important sugarcane feedstock quality traits, namely fiber (FIB) and sucrose (PC) content, in a commercial sugarcane (*Saccharum* spp.) population from the sugarcane genetic breeding program of the Universidade Federal de Viçosa (PMGCA-UFV), genotyped for single nucleotide polymorphisms (SNPs) and screened using NIR spectroscopy. We compared three modelling strategies: 1) genome-based prediction, 2) NIR-based prediction, and 3) the integration of SNP markers and NIR wavenumber variables as predictors.

## Material and methods

### Plant material

In this study, we evaluated 385 clones derived from an originally seedling population of 98 half-sib families. The seedling population, in which each plant is a single genotype, was the result of crosses made at the Serra do Ouro Flowering and Breeding Station, municipality of Murici, Alagoas State, Brazil (09°13’ S, 35°50’ W, 450 m altitude). After processing, seeds were sent to the Sugarcane Genetic Breeding Research Station (CECA) of the Universidade Federal de Viçosa, municipality of Oratórios, Minas Gerais State, Brazil (20°25’ S, 42°48’ W, 494 m altitude) and germinated in a nursery house. Subsequently, seedlings obtained from each family were transplanted to the field and evaluated in first (plant cane) and second (ratoon) crops based on desirable traits in first (T1) and second (T2) clonal trial phases [23].

### Experimental design

An augmented block design was initiated in May 2016 at the CECA municipality of Oratórios, Minas Gerais State, Brazil (20°25’ S, 42°48’ W, 494 m altitude). The checks and released cultivars RB867515, RB966928, and RB92579 were included once in each block, and regular treatments were arranged in 21 blocks [24]. In total, 18 blocks contained 24 plots; one block contained 16 plots, and the remaining two blocks contained 12 plots. The experimental plots consisted of double-row 3 m long furrows × 1.4 m between rows and clones were cultivated following standard agronomic protocols regarding fertilization, weed control, and pest management [25]. Buffer rows of a released cultivars encompassed the whole experiment area.

### Phenotypic data

The clones were evaluated in the first ratoon (second crop), 26 months after planting. The method employed to estimate the percentage of FIB content followed recommendations of the CONSECANA manual [26]. Sugarcane breeding programs in Brazil routinely apply these protocols. Harvest and quality analyses were performed in July 2018. To obtain a representative set of the samples, ten randomly selected stalks from each of the double-row plots were cut at ground level with a machete. Green tops, clinging leaves, and leaf sheaths were removed before stalks were bundled and weighted using a dynamometer (S1 Fig). These ten randomly selected stalks in each plot were shredded. A subsample of 500 g from the shredded stalks was collected and pressed with a hydraulic press. After pressing, the remainder fiber cake was collected and taken to the laboratory. We obtained the BRIX% (percentage of soluble solids) and POL% (percentage of sucrose) values from the juice. The BRIX was obtained with a refractometer (HI96801 Model, Hanna® instruments, Woonsocket, USA), while POL was obtained by polarimetry using a saccharimeter (SDA2500 Model, Acatec, Brazil) after clarifying the solution with lead acetate. The remainder fiber cake was weighed (*WC*) and used to derive fiber content [27]:

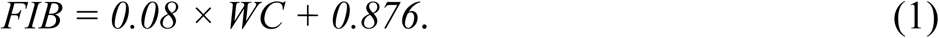

The apparent percentage of sucrose in sugarcane (PC%) was derived based on POL% and FIB as follows:

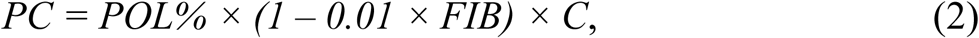

where *C* is the coefficient to convert sucrose of juice into sucrose of cane calculated using the formula *C = 1.0313 – 0.00575 × FIB*. The final values were all expressed as the total fresh biomass basis (500 g of shredded stalks).

### Sample preparation and NIR spectra acquisition

Another subsample of 100 g from the shredded stalks was collected and immediately taken to dry in a forced-air circulating oven at 50°C for 24 h or until a constant mass was reached (S1 Fig). Dried samples were then ground, packaged in a plastic zip bag, and stored. The NIR spectra of samples were measured in indoor-conditions at room temperature of 21 °C. The instrument used was a Fourier transform near-infrared (FT-NIR) spectrometer set (Antaris™ II Model, Thermo Scientific Inc., USA), under the following operating conditions: 4 cm^−1^ resolution in an investigated wavenumber range of 10000 to 4000 cm^−1^ and reflectance mode as log (1/R), where R is the measured reflectance. Samples were placed into a powder sampling cup accessory and arranged into the instrument window. At each scan, the accessory was moved to cover different positions of the sample, totaling six positions. A single scan measure was the average result of 32 scans. For each sample, a total of 192 scans were made and then averaged, representing the final spectrum. The final NIR matrix used in the subsequent analyses had a dimension of 385 rows and 3,112 columns.

### DNA isolation, sequencing, and genotyping data

Sugarcane DNA samples were isolated using DNeasy Plant Mini Kit (QIAGEN, Hilden, Germany) and sent to RAPiD Genomics (Gainesville, Florida, USA) for the construction of probes, sequencing, and identification of molecular markers. Samples were genotyped using single-dose SNP markers based on the Capture-Seq technology (https://www.rapid-genomics.com). Raw sequence reads were mapped, called, and filtered. Reads were anchored to a monoploid reference genome of sugarcane (*Saccharum spp*.) [28] using the BWA-MEM algorithm of the BWA version 0.7.17 [29], and a flag identifying the respective sugarcane genotype was added to each mapping file. The mapping files were processed using SortSam, MarkDuplicates, and BuildBamIndex tools of Picard version 2.18.27 (https://github.com/broadinstitute/picard/). Variants were called using FreeBayes version 1.2.0 (https://github.com/ekg/freebayes) with a minimum mapping quality of 20 (probability of miscalling), minimum base quality of 20 (SNPs with missing data higher than 20% were eliminated), and minimum coverage (how many times a fragment was sequenced) of 20 reads at every position in the reference genome. Thereafter, the SNP marker matrix was coded counting the occurrence of the reference allele A. Thus, considering the genotypes AA, Aa, and aa, the matrix entries would be 2 (homozygosity for the reference allele), 1 (heterozygosity with one reference and one alternative allele), and 0 (homozygosity for the alternative allele), respectively. Further, markers with minor allele frequency lower than 5% were eliminated. Lastly, missing variants were imputed from a binomial distribution density function using the frequency of the non-missing variants. A total of 124,307 SNPs was retained for further analyses.

### Statistical Analysis

A two-stage analysis was employed. In the first step, we run a mixed model equation with variance components estimated by REML using the SELEGEN-REML/BLUP software [30]. We considered the model

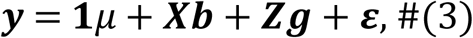

where **y** is the vector of phenotypes; **1** is a vector of 1s; µ is the overall mean; **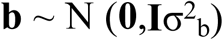** is the vector of random block effects; 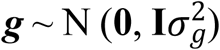 is the vector of random genetic effects, and **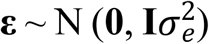** is the vector of residuals. The capital letters **X** and **Z** represent the incidence matrices of the respective random effects. In the second step, the best linear unbiased predictors (BLUPs) of each trait were used as dependent variables using three prediction models (M1, M2, and M3) to evaluate the prediction accuracy for FIB and PC.

The first (M1) was a single-trait model using only markers which takes the following form:

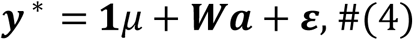

where ***y***^*^ is the vector of adjusted phenotypic values (BLUPs) for FIB or PC, **1** is a vector of 1s; µ is the overall mean, **W** is the matrix with SNP markers for each individual, ***a*** is the corresponding vector of marker effects, and **ε** is the vector of residuals.

The second (M2) was a single-trait model using NIR wavenumber variables as predictors:

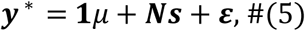

where ***N*** is the matrix with the spectrum for each individual along the wavelength, and ***s*** is the corresponding vector of wavelength effects. It is a commonplace to apply mathematical transformations to the NIR matrix before analysis to increase the signal to noise ratio [31]. More details can be found elsewhere [32]. We tested different combinations of pre-processing techniques. The pre-processing combination that yielded the best results was Savitzky-Golay smoothing (SGS) (window: 5; polynomial order: 2) followed by multiplicative scatter correction (MSC) and mean centering (MC) for FIB, whereas for PC the best combination was SGS (window: 5; polynomial order: 2) and MC (Fig 1).

**Fig 1.**
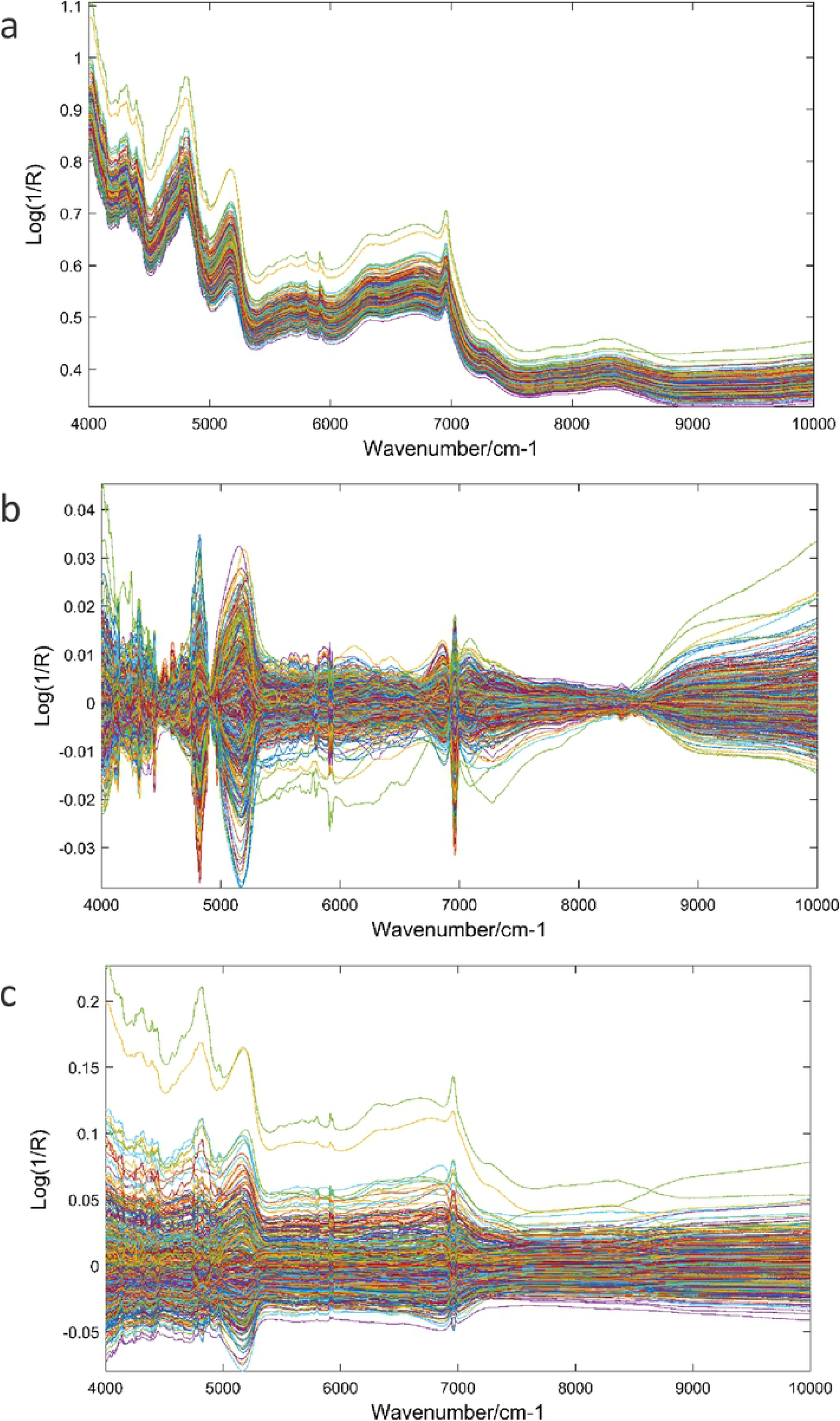
Sugarcane samples raw spectra (a) and pre-processed spectra for fiber content (b), and sucrose content (c).

In the third model (M3), we combined SNP markers and NIR wavenumber variables as predictors fitting the following linear model:

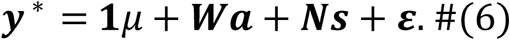

In equation [6], the pre-processing combination of the NIR matrix that best contributed with the SNP matrix for maximizing the prediction accuracy were SGS (window: 5; polynomial order: 2), MSC and MC for FIB, and SGS (window: 5; polynomial order: 2), 1° derivative, MSC and MC for PC. The incidence matrices ***N*** and ***W*** were scaled (centered and standardized) in all prediction models prior to the analyses.

### Regression models

The models were tested using two regression methods: The BayesB and partial least squares (PLS). BayesB is a hierarchical Bayesian approach that performs variable selection [3]. We used a multi-layer BayesB by assigning different independent priors for SNP markers and NIR wavenumbers in M3. PLS regression is a dimension reduction method and fundamentally transforms the original collinear predictors into non-correlated variables [33]. In PLS regression, the algorithm identifies the principal components (latent variables) that best describe the data in terms of variance, and it does so by constructing linear combinations of all predictors. Furthermore, unlike other dimension reduction models such as principal component regression, the fitting procedure of PLS involves finding the latent variables that maximize the covariance between the predictors and phenotypes while minimizing the error [34,35].

The BayesB analyses were carried out using the BGLR package (Pérez; et al., 2014). We run the BayesB for 25,000 samples, with the first 10,000 being discarded (burn-in) with a thinning interval of 10. The PLS regression was performed using the mixOmics package [37]. Both methods were implemented in R [38].

### Accuracy of predictions

The prediction accuracy of models M1, M2, and M3 was evaluated by random sub-sampling cross-validation repeated 20 times. The models are fitted using the data of the training set observations and tested to predict unknown samples of the validation set. In this study, the training set contained 80% of the samples (308 clones) and the validation set included the remainder of 20% (77 clones). At each time, the algorithm randomly selected a different subset of observations assigned to the training and validation sets. The results were compared by computing the mean Pearson correlation coefficient between observed and predicted genotypic values. A schematic diagram summarizing the whole experimental preparation and processing is depicted in Fig 2.

**Fig 2.**
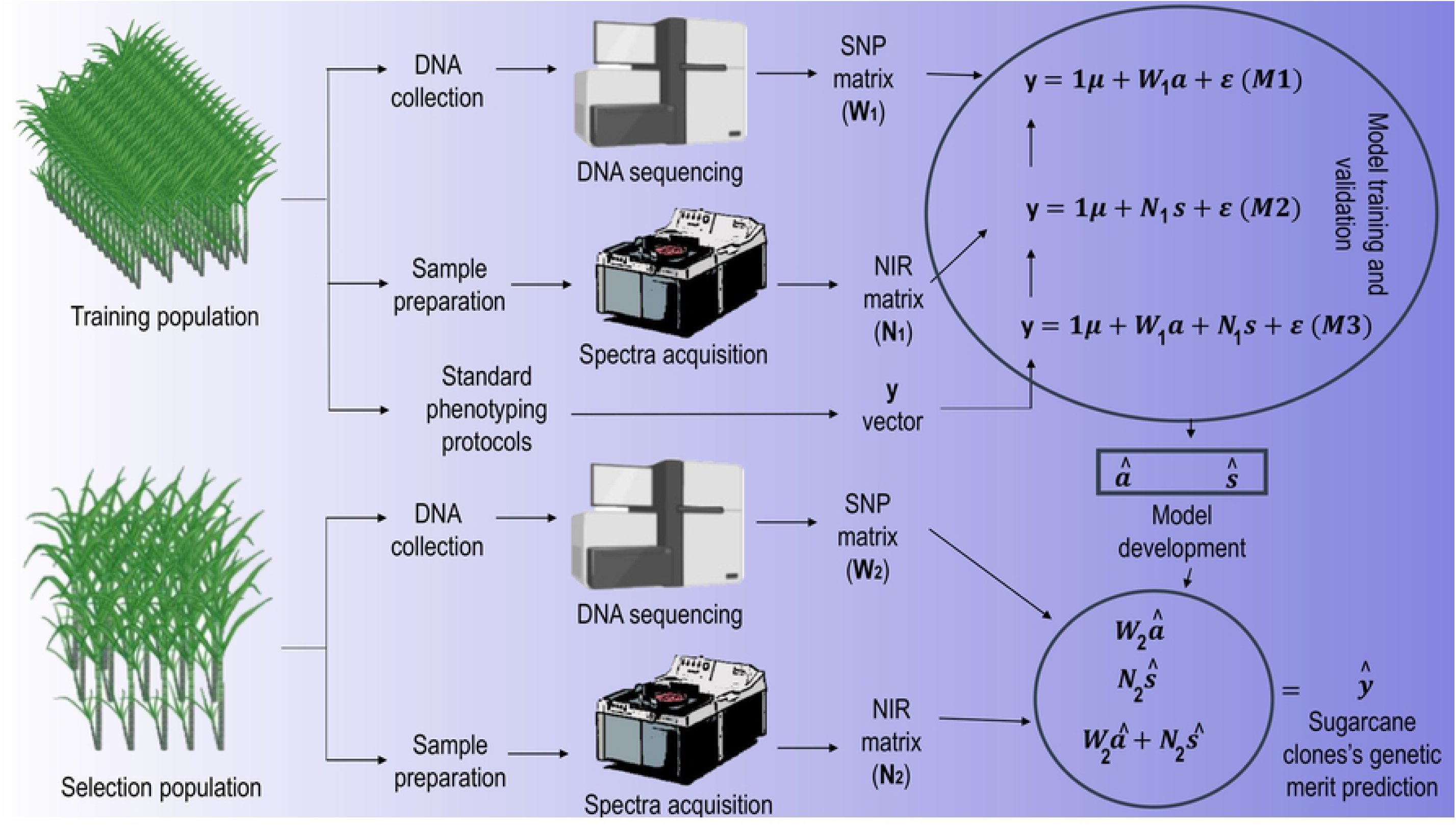
Schematic diagram of the experimental procedure.

## Results

### Phenotypic data

Fig 3 shows the correlation analysis using the BLUPs of FIB and PC. The data followed a Gaussian distribution curve. The two traits evaluated are negatively correlated (r = -0.22; p < 0.001).

**Fig 3.**
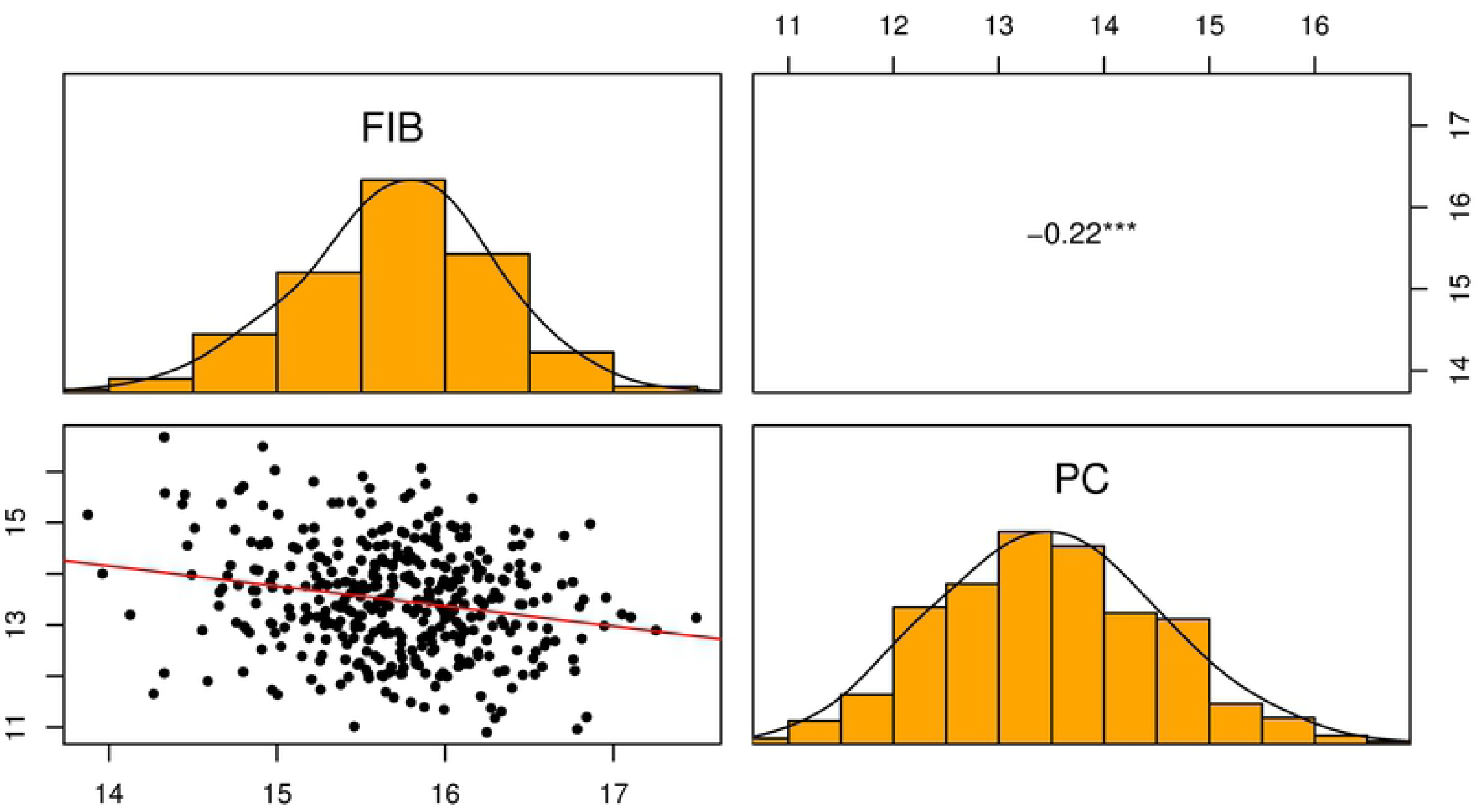
Scatter plot, histogram, and correlation of BLUPs of FIB and PC for 385 sugarcane clones.

The variance component estimates calculated using the REML/BLUP procedure were used to derive genetic and environmental parameters. Significant values (p < 0.01) of genotypic variance 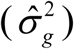 were observed from the deviance analysis for FIB and PC (Table 1). The estimates of individual broad-sense heritability (*h*^2^) for FIB were considerably high. In contrast, the *h*^2^ for PC was low. The heritability values obtained might have been the result of environment variance and the choice of the experimental design.

**Table 1.**
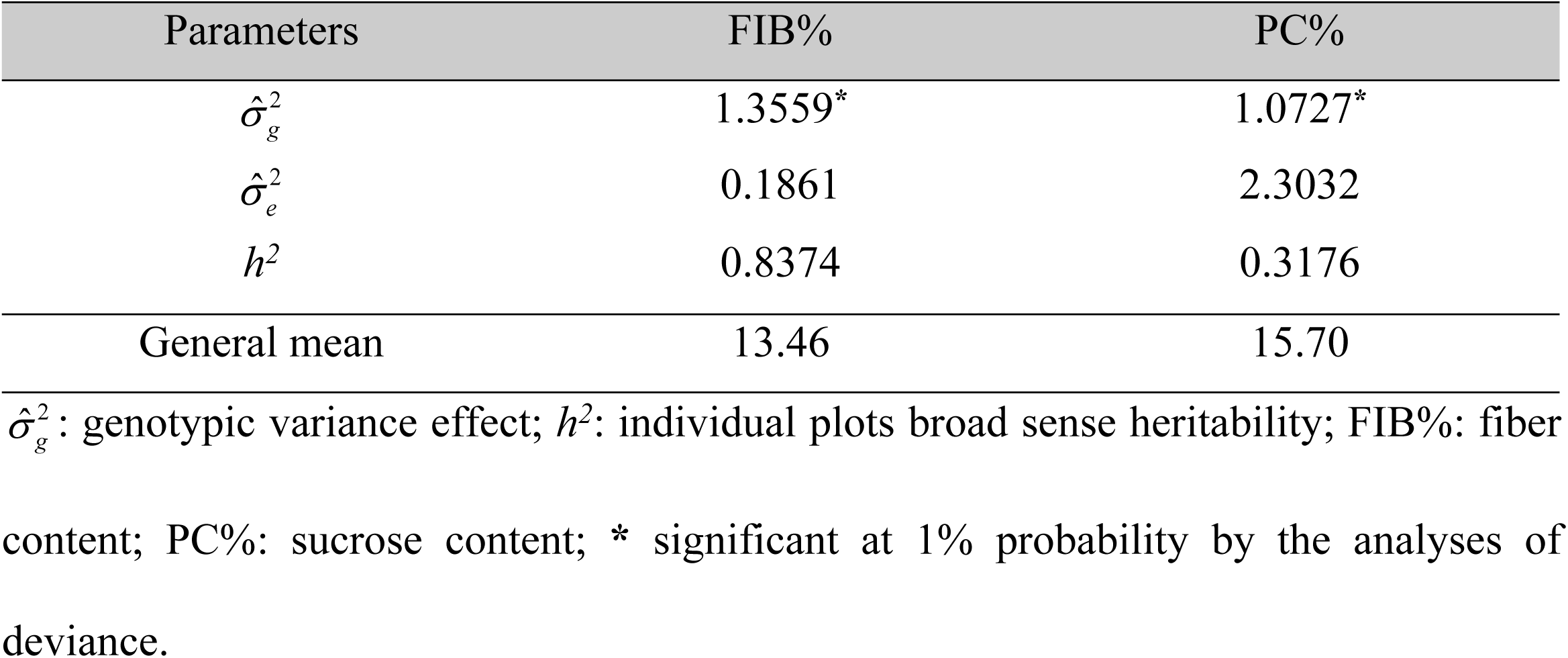
Genetic and environmental parameter estimates of the 385 sugarcane clones evaluated.

### Prediction models

The BayesB results from cross-validation are presented in Fig 4. We found that M1 resulted in the lowest prediction accuracy for FIB and PC. Additionally, M1 models showed the largest cross-validation uncertainty. The highest prediction accuracy of M2 was obtained for FIB (0.6138), followed by PC (0.5447). The combination of SNP markers and NIR spectra in M3 models yielded an increase in the predictive ability for PC (0.5860) and a marginal improvement for FIB (0.6231) in comparison to the models fitted using only NIR spectra (M2) as predictor variables.

**Fig 4.**
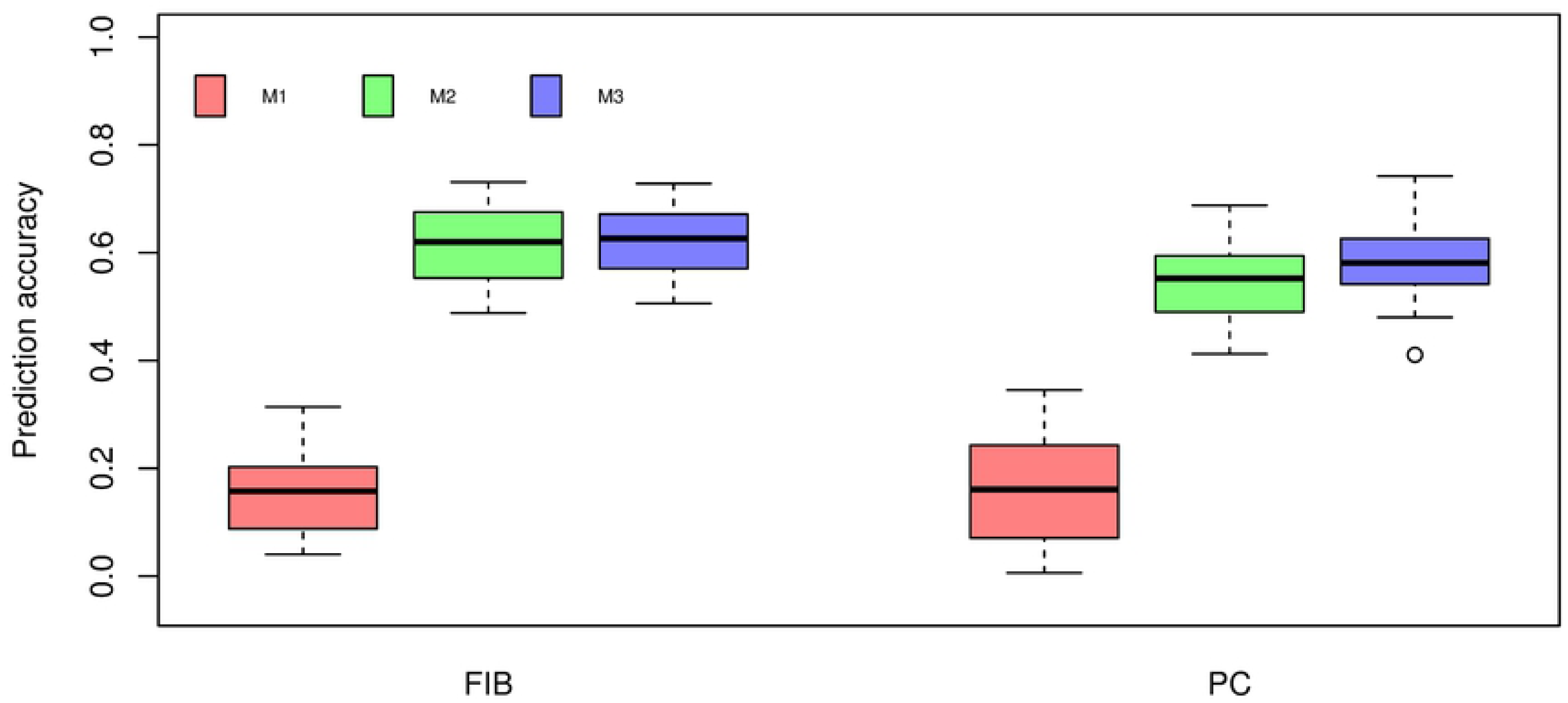
Box-plot of cross-validation prediction accuracy of fiber content (FIB) and sucrose content (PC) using BayesB under three different prediction models. M1: markers. M2: near-infrared spectra. M3: the combination of markers and near-infrared spectra.

The cross-validation results using PLS regression are shown in Fig 5. Similar to BayesB, M1 models fitted using PLS resulted in the lowest prediction accuracies for both traits. The M2 model for FIB presented the highest prediction accuracy across prediction models (0.3917). Considering M3 models, we observed a small increase for PC (0.3942) than that of the M2 model using NIR spectra alone (0.3673). In contrast, no improvement in prediction accuracy was observed for FIB.

**Fig 5.**
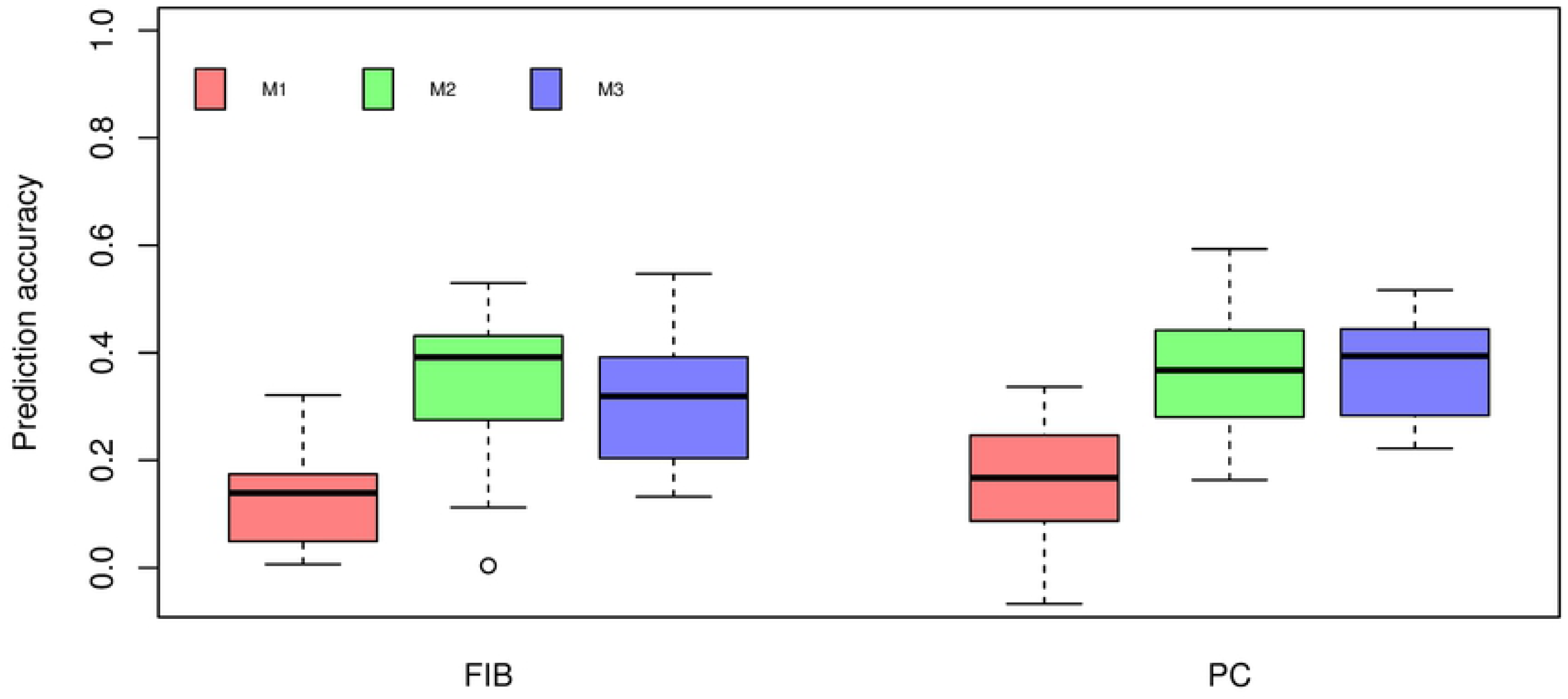
Box-plot of cross-validation prediction accuracy of fiber content (FIB) and sucrose content (PC) using PLS under three different prediction models. M1: markers. M2: near-infrared spectra. M3: the combination of markers and near-infrared spectra.

## Discussion

In the present study, we explored different strategies to incorporate genomic and HTP derived information from NIR spectroscopy into a sugarcane genetic breeding program aiming to improve prediction accuracies. Jannink et al. (2010) [39] reported the similarity of NIR spectroscopy and GS approaches, as they inherently share the same purposes and statistical analysis challenges. For instance, the application of NIR spectroscopy aims to replace demanding and expensive laboratory protocols by developing statistical prediction models using high-dimensional input variables. Likewise, GS is intended to utilizes multivariate statistic methods to build prediction models that associate difficult to record plant phenotypes with easy to measure variables (e.g., SNPs) [40]. Moreover, the existing statistical methods capable of coping with the challenges associated with high-dimensional statistical analysis are used across research fields [22,41–44].

Along with the integration of genomic and spectroscopy datasets, we also considered their application as single predictors in the statistical models and evaluated the impact in the prediction accuracy of two sugarcane feedstock quality traits using a commercial sugarcane population of 385 individuals and tested two regression methods. We assessed the performance of PLS regression, which is arguably the most employed method when dealing with NIR datasets [45] and BayesB for the GS models. For all models, we employed repeated random sub-sampling cross-validation to assess the accuracy of prediction models.

### Phenotypic data

The results we found for genotypic effects suggest the presence of genetic variance to be exploited for selection purposes. These results are in agreement with the study of Baffa et al. (2014) [46] and Wang et al. (2008) [47] for commercial sugarcane populations.

The estimated broad-sense heritability values indicate that the selection of clones for these trait based on phenotypic values would be effective since environment variance has shown little impact [48]. Ramos et al. (2017) [49] found similar values of *h*^2^ for FIB and PC.

The negative correlation observed between FIB and PC support the dynamic of carbon partitioning and the antagonistic metabolic pathway of fiber and sucrose synthesis in sugarcane related in other studies, which suggest the trend of PC being negatively impacted by the increase of FIB [50,51].

### Prediction models

Overall, models fitted using BayesB were most predictive than PLS models for both traits, with accuracy estimates ranging from 0.5860 to 0.6231 (Figure 5). This result is in agreement with the study of Ferragina et al., (2015) [44], in which Bayesian models outperformed PLS regression for NIR-based prediction of dairy traits. Additionally, Solberg et al. (2009) [52] reported that the prediction accuracy of genome-wide breeding values from PLS regression was lower compared to that of BayesB.

The GS models (M1) showed poor predictive ability and only explained a small portion of the phenotypic variation of the two traits evaluated. The predictive ability of GS models observed was lower than that found by Gouy et al., (2013) [42]. The authors reported accuracies for within cross-validation panels considering bagasse content (an equivalent measure of the FIB trait we used herein) and BRIX, which is highly correlated with PC [53], of up to 0.5 and 0.62, respectively. However, they used a DArT low-density molecular marker panel, whereas we used a high-density SNP panel.

Numerous factors can affect the results of GS, for instance, marker density and the amount of existing linkage disequilibrium (LD). Jannink et al. (2010) [39] reported that the size and the genetic relatedness between training and validation population genotypes is a paramount aspect. The genome structure of the population in which we want to infer breeding values should be similar to the one we used to train our model. In the present study, no training population optimization was performed since the population used was composed of clones coming from an advanced stage of selection. Therefore, it can be argued that this might be the reason for the lower prediction accuracies we obtained [54].

The GS models fitted in this study only considered additive effects. However, Zeni Neto et al. (2013) [55] reported that additive and non-additive genetic effects are equally crucial for the determination of complex traits in sugarcane. Hence, the inclusion of non-additive effects in GS models for sugarcane and other clonally propagated species may improve prediction accuracies [56]. Results obtained by Denis et al. (2013) [57] in a simulation study, and de Almeida Filho et al. (2014) [58] using data from a full-sib population of loblolly pine (*Pinus taeda* L.) indicate the increase of prediction accuracy when accounting for dominance effects. Also, the incorporation of pedigree information is suggested to improve accuracies compared to the sole use of molecular markers [59].

The increase in coverage is another highlighted aspect of GS [60,61]. However, Sousa et al. (2019) [62] applied GS in polyploid hybrids of *Coffea* spp. and suggested that once the optimal number of SNPs is reached, a plateau in terms of the increase in selective accuracy is observed, and then decreases. Yang et al. (2017) [63] reported that for sugarcane and other crops with a complex genome, the quality of sequencing might be more important than a large number of SNPs and, therefore, sequencing depth is paramount to filter low-quality sequence reads.

These pieces of evidence suggest that the most significant bottleneck of the application of GS in sugarcane and other complex genome species can be addressed in the sequencing and sequencig data processing steps, which is a relevant issue that is currently receiving many research efforts [64–66].

The M3 modelling strategy, we tested in this study aimed to increase the prediction accuracy by incorporating NIR wavenumbers and markers to predict sugarcane clones’ genetic merit. We expected that the combination of predictors would bring synergy and thus, improve model performance. However, the combination of predictors in a single model showed no significant improvement in comparison to the use of NIR wavenumber variables alone. Seemingly, most of the variation of the two evaluated traits that were captured by the model that combined NIR and SNP predictors came from the NIR spectra.

Crain et al. (2018) [22] investigated a similar strategy we proposed herein and found the same trend. The authors evaluated the effect of including HTP data into GS models in different stressed environmental conditions. The results of their study revealed that models including HTP derived information as single predictors contained most of the predictive ability when compared to models with markers alone in one of the evaluated scenarios, which is consistent with the results we observed.

Cost and efficiency are key drivers to determine the adoption of GS and HTP in a breeding program. One of the main interests is to optimize the selection process by reducing breeding cycles, saving time, and reducing costs by performing the early selection of candidate elite genotypes. Currently, at the PMGCA, clonal performance is assessed by field-based collection of data through multiple harvests and at different locations [67]. Clones are evaluated in plant cane, and the selection is made in the first ratoon. After hybridizations, the first stage of clonal selection at the PMGCA is referred to as the T1 phase [68]. At the T1 phase, seedlings are transferred to the field, and mass selection is performed. Hence, the selection accuracy is very low, especially regarding low heritability traits [4]. Consequently, at T1, the replacement of visual appraisal phenotypic selection by high-throughput scientifically gathered data is highly desirable, as it could improve selection accuracy [69].

Rincent et al., (2018) [70] have proposed an approach in which relationship matrices are derived from NIR spectra data and compared the efficiency of predictions with standard GS models considering markers. The results found by these authors suggest that models developed using NIR data can outperform GS models. Likewise, we observed with our dataset that NIR models alone provided better results than GS models. Other works have also demonstrated the feasibility of using HTP data to improve predictions. For instance, Krause et al., (2019) [69] used hyperspectral derived relationship matrices to model genotype × environment interactions. Also, the advent of portable NIR instruments is heralded to become a useful low-cost tool to assist plant breeders [71]. Consequently, it may provide better results over genotyping for high-density SNP panels of candidate sugarcane clones, as this approach also demands time and resources, and at the moment, it lacks the desired effectiveness.

## Conclusion

Our results show that GS models had the lowest prediction accuracies and the integration of NIR wavenumber variables and SNP markers as predictor variables to predict sugarcane feedstock quality traits have not demonstrated significant differences when compared to the models including only NIR wavenumber variables.

NIR wavenumber variables alone achieved high prediction accuracies for the two traits assessed in this study, which suggests that NIR spectroscopy could be used as an efficient tool to maximize resources and for predicting the genotypic value of clones in a sugarcane breeding program.

## Acknowledgments

The authors thank Professor Dr. Luis Antônio dos Santos Dias for providing the NIR instrument used in this study. Also, we acknowledge Professor Dr. Reinaldo Francisco Teófilo and Dr. Jussara Valente Roque for the helpful assistance regarding NIR spectra collection and downstream analyses. Finally, we acknowledge the numerous co-operators from the Sugarcane Genetic Breeding Research Station (CECA-Minas Gerais State, Brazil) who helped carrying out field trials and to collect phenotypic data.

## Supporting information

**S1 Fig. Overview of data acquisition.**A: stalks being harvested from double-row plots; B: stationary forage chopper machine used to shred stalks; C: hydraulic press used to extract the fiber cake and juice samples; D: fiber cake being weighted; E: samples being dried at a forced-air circulating oven; F: dried ground samples placed onto the NIR instrument window; G: saccharimeter instrument; H: sample spectrum displayed on the computer screen.

